# The Use of iPSC-Derived Cardiomyocytes and Optical Mapping for Erythromycin Arrhythmogenicity Testing

**DOI:** 10.1101/574012

**Authors:** A.D. Podgurskaya, V.A. Tsvelaya, M.M. Slotvitsky, E.V. Dementyeva, K.R. Valetdinova, K.I. Agladze

## Abstract

Erythromycin is an antibiotic that prolongs the QT-interval and causes Torsade de Pointes (TdP) by blocking the rapid delayed rectifying potassium current (I_Kr_) without affecting either the slow delayed rectifying potassium current (I_Ks_) or inward rectifying potassium current (I_K1_). Erythromycin exerts this effect in the range of 1.5 μM–100 μM. However, the mechanism of action underlying its cardiotoxic effect and its role in the induction of arrhythmias, especially in multicellular cardiac experimental models, remain unclear. In this study the re-entry formation, conduction velocity, and maximum capture rate were investigated in a monolayer of human induced pluripotent stem cell (iPSC)-derived cardiomyocytes from a healthy donor and in a neonatal rat ventricular myocyte (NRVM) monolayer using the optical mapping method under erythromycin concentrations of 15, 30, and 45 μM. In the monolayer of human iPSC-derived cardiomyocytes, the conduction velocity (CV) varied up to 12±9% at concentrations of 15–45 μM as compared with that of the control, whereas the maximum capture rate (MCR) declined substantially up to 28±12% (*p* < 0.05). In contrast, the tests on the NRVM monolayer showed no significant effect on the MCR. The results of the arrhythmogenicity test provided evidence for a “window” of concentrations of the drug (15 to 30 μM) at which the probability of re-entry increased.

## Introduction

As reported in the cardiology literature, erythromycin is a macrolide antibiotic that prolongs the QT-interval and causes Torsade de Pointes (TdP) [1,2]. Previous research demonstrated an association between erythromycin, an increased risk of sudden cardiac death and ventricular arrhythmias [3]. This drug prolongs the repolarization phase of an action potential [4] by blocking the component I_Kr_ (rapid delayed rectifying potassium current) of the current through voltage-gated potassium channels [5–8]. The component I_Ks_ (slow delayed rectifying potassium current) and the inward rectifying potassium current I_K1_ are unaffected by erythromycin [6].

The therapeutic blood plasma concentration of erythromycin is 3–8 μM [9], although the effective concentration may be as high as 30 μM [2]. In experiments *in vitro*, erythromycin had a significant effect on the field potential duration (FPD) in human embryonic stem cell-derived cardiomyocytes at concentrations of 1.5–16 μM, but it did not induce proarrhythmic effects [10].

At physiological temperature, erythromycin blocked hERG channels, with an IC50 (the half maximal inhibitory concentration) at a concentration of 40 μM[11]. The same effect was achieved by concentrations of erythromycin up to 100 μM under physiological temperature conditions but not under room temperature (24° C conditions [12]. Only a few studies have focused on the mechanism of action of erythromycin in multicellular cardiac experimental models [2, 13–15], and the role of this drug in the occurrence of arrhythmias remains unclear. Furthermore, information is lacking on the influence of varying erythromycin concentrations on excitation wave propagation.

In this study, we investigated the role of erythromycin in re-entry formation in a monolayer of human iPSC-derived cardiomyocytes from a healthy donor and in a neonatal rat ventricular myocyte (NRVM) monolayer. The conduction velocity (CV) and maximum capture rate (MCR, the maximum rate at which each stimulus was followed by a response) were evaluated using the optical mapping method at erythromycin concentrations of 15, 30, and 45 μM. The dose-dependence arrhythmogenicity of erythromycin on a standard linear obstacle was determined. The experiments also demonstrated the mechanism of re-entry formation and the probability of its appearance, which depended on the stimulation frequency and the concentration of erythromycin.

## Materials and Methods

### Preparation of samples

Pre-fired cover glasses (13 mm in diameter and 22 mm in diameter) were placed in the sterile 24-well culture plates or in Petri dishes. Further, the glasses were burned with ultraviolet for 30 minutes. To improve the adhesion of NRVMs, human fibronectin (Imtek) was used. After application of fibronectin to each cover glass at a concentration of 20 μg/ml, the Petri dishes were transferred to an incubator (37 °C, 5% CO_2_) for 12 hours. iPSCs were cultured in the Essential 8™ Medium on a Geltrex LDEV-Free hESC-Qualified Reduced Growth Factor Basement Membrane Matrix (ThermoFisher Scientific) after transferring from the feeder layer. For experiments, iPSCs were seated in the wells of 24-well sterile plates covered with Geltrex. The differentiation protocol started when the 80% of confluence monolayer was formed on day 3-4 after plating.

All studies were conducted by the National Institutes of Health Guide for the Care and Use of Laboratory Animals (NIH publications No. 8023, revised 1978) and approved by the Moscow Institute of Physics and Technology Life Science Center Provisional Animal Care and Research Procedures Committee, Protocol \#A2-2012-09-02.

### Fibroblast derivation and reprogramming to the pluripotent state

Dermal fibroblasts were isolated from a skin biopsy of healthy donor. Fibroblasts were cultivated in the medium contained DMEM:F12 1:1, 15% fetal bovine serum, 1×GlutaMAX™ Supplement, 1×Penicillin-Streptomycin. Fibroblasts were nucleofected with episomal vectors expressed *OCT4*, *SOX2*, *KLF4*, *L-MYC*, and *LIN28* (Addgene IDs #41855-41858, #41813-41814) on a Lonza Nucleofector 2b. The cells were plated on a surface coated with Geltrex LDEV-Free hESC-Qualified Reduced Growth Factor Basement Membrane Matrix. Reprogramming to the pluripotent state was carried out as described in the protocol to the Epi5™ Episomal iPSC Reprogramming Kit (https://tools.thermofisher.com/content/sfs/manuals/epi5_episomal_ipsc_reprogramming_man.pdf). Colonies similar to those of human pluripotent stem cells appeared beginning from day 15 after nucleofection. The colonies were transferred with capillary to a feeder layer of mitotically inactivated mouse fibroblasts and cultivated up to generating stable iPSC lines in the medium contained 85% Knockout DMEM, 15% Knockout Serum Replacement, 1×GlutaMAXTM Supplement, 1×Penicillin-Streptomycin, 1×Non-Essential Aminoacids, 0.05 mM β ercaptoethanol (Sigma-Aldrich), 10 ng/ml basic Fibroblast Growth Factor (Biolegend). All reagents were from ThermoFisher Scientific if another supplier is not stated.

### Characterization of iPSC lines

Analysis of expression of alkaline phosphatase was performed as described previously [14]. Spontaneous differentiation of the cell lines was performed through embryoid bodies formation [16]. 14-day embryoid bodies were treated with 0.25% trypsin. Cell suspension was plated on 4-well plates and cultivated for 7 days. Immunofluorescent analysis of iPSCs and their differentiated derivatives was carried out as described below.

### Immunofluorescent staining

Cells were fixed for 10 min in 4% paraformaldehyde, permeabilized for 10 min in 0.4% Triton-X100 or 96% ethanol (for TRA-1-60 and TRA-1-81). Immunostaining with antibodies to SSEA-4 was done without permeabilization. Cells were further incubated for 30 min in blocking buffer (1% bovine serum albumin in Phosphate Buffered Saline, PBS), overnight at 4ºC with primary antibodies and for 1 h at room temperature with secondary antibodies. Cells were washed twice for 15 min in PBS. Nuclei were stained with DAPI. Samples were analyzed on a Nikon Eclipse Ti-E.

Primary antibodies and working dilutions — TRA-1-60 (Abcam, ab16288) 1:200, TRA-1-81 (Abcam, ab16289) 1:200, SSEA-4 (Abcam, ab16287) 1:50, NANOG (ReproCELL, RCAB003P) 1:200, OCT4 (Santa Cruz, sc-5279) 1:200, beta-III-tubulin (Abcam, ab7751) 1:100, cytokeratin 18 (Millipore, MAB3234) 1:200, smooth muscle α-actin (DAKO, M0851) 1:100, CD90 (Millipore, MAB1406) 1:100, cardiac troponin T (Abcam, ab8295) 1:100, β-myosin light chain 2 (Abcam, ab79935) 1:100, β-myosin heavy chain (Abcam, ab15) 1:100, NKX2.5 (Santa Cruz, sc-14033) 1:100. Secondary antibodies (ThermoFisher Scientific, working dilution - 1:400) - Alexa Fluor 568 goat anti-mouse IgG (H+L) highly cross adsorbed (A11031), Alexa Fluor 488 goat anti-mouse IgG (H+L) highly cross adsorbed (A11029), Alexa Fluor 488 goat anti-mouse IgG1 (A21121), Alexa Fluor 568 goat anti-rabbit IgG (H+L) (A11011), Alexa Fluor 488 goat anti-rabbit IgG (H+L) (A11008), Alexa Fluor 488 goat anti-rat IgG (H+L) cross adsorbed (A11006).

### Directed differentiation of iPSCs into cardiomyocytes

iPSCs were cultivated for several passages under feeder-free conditions (in the Essential 8™ Medium on Geltrex LDEV-Free hESC-Qualified Reduced Growth Factor Basement Membrane Matrix (ThermoFisher Scientific). 3-4 days before differentiation, iPSCs were plated on 24-well plates. Directed differentiation of iPSCs into cardiomyocytes was triggered at 80-90% cell density by adding the RPMI 1640 medium (Lonza) contained B27 supplement minus insulin (Thermo Fisher Scientific) and 8 μM CHIR99021 (Sigma-Aldrich) for 48 h. Other differentiation steps were carried out as described in Supplementary materials to [15]. Starting from day 9 of differentiation, the first cell contractions were observed. Optical mapping occurred when the culture reached 50 days.

For immunofluorescent staining, differentiated cells were disaggregated with TrypLE™ Express (ThermoFisher Scientific) and plated on slides coated with 0.1% gelatin in the medium contained 80% RPMI 1640, 20% fetal bovine serum, and 10 μM Y-27632 (StemRD).

### Preparation of neonatal rat cardiomyocytes

Cardiac cells were isolated according to the Worthington protocol (http://www.worthington-biochem.com/NCIS/default.html). In this work the hearts of rats Rattus norvegicus, Sprague Dawley breed aged 1–4 days were used. Seating of the cells occurred for optical mapping at a concentration of 300 thousand/cm^2^ drop on the prepared glass. After plating the cells, the samples were placed in an incubator (37 °C, 5% CO_2_) for 1–2 hours. Then DMEM supplemented with 10% FBS, 2 mM of L-glutamine, and 100 U/ml of penicillin/streptomycin was added to each sample and the samples were returned to the incubator for 24 hours. The next day there was a change of medium on DMEM described above with 5% FBS. After 3–4 days of cultivation, the cells were observed in a light microscope for the formation of a confluent monolayer and the formation of contractile syncytium, which served as an indicator for the possibility of optical mapping of this sample.

### Optical mapping of neonatal rat cardiomyocytes

After 3–4 days of culturing the monolayer of neonatal rat cardiomyocytes, the fluorescent calcium-dependent fluo-4 dye (https://www.thermofisher.com/order/catalog/product/F14201) in concentration 4 μg/ml was applied to the samples for 35–40 minutes without access to light. Then the dye solution was exchanged with a fresh heated solution of Tyrode salts (pH = 7.4). All experiments were carried out at a temperature of 37 °C. Optical mapping was carried out on the installation consisted of a high-speed video camera (Andor IXon3, Andor Technologies), a mercury lamp (Olympus U-RFL-T), an optical microscope (Olympus MVX10), a filter cube (Olympus U-M49002XL, (Vellemann, PCGU-1000), a platinum point electrode and a reference circular electrode. The electrodes were used for point stimulation of the cardiac tissue.

Each sample was checked for spontaneous activity. The amplitude of the pulses was not higher than 8V. The video was shot at a speed of 68–130 FPS (frames per second) and a spatial resolution of ~ 0.03 mm / pixel.

### Data processing

All videos from optical mapping were processed in the ImageJ program. The activation maps were built using the Wolfram Mathematica program. The image processing from the confocal microscope was performed in the Zen program corresponding to the microscope software.

### Optical mapping of human iPSC-derived cardiomyocytes

Optical mapping of human iPSC-derived cardiomyocytes occurred 50 days after the start of differentiation protocol. The sample incubated in a sterile medium at 37 ° C with a fluor-4 dye in concentration 4 μg/ml for 30 min. Then the dye solution was exchanged with a sterile Tyrode’s solution. All experiments were carried out at a temperature of 37 °C.

### Protocol of optical mapping

After capturing the CV and frequency controls, the solution of erythromycin (JSC Sintez) in Tyrode’s was added to the cell culture at different concentrations: 15 μM, 30 μM, 45 μM. For each concentration of the substance, the incubation time was more than 5 minutes. The video was recorded after the effect of the substance did not change for 15 minutes.

### Standard linear obstacle

After adding Tyrode’s to the sample, a standard linear obstacle was made on the tissue culture perpendicular to the direction of excitation wave propagation using a thin blade. The obstacle width was ≤100 μm. The configuration of the propagating wave front under normal conditions and under various (15–45 μM) concentrations of erythromycin in the presence of a standard linear obstacle was evaluated to indicate re-entry formation. The dose-dependence arrhythmogenicity of erythromycin was determined.

### Statistics

All experimental numerical data are presented as the mean ± SD unless otherwise specified. The statistical significance was determined using a one-way ANOVA followed by Fisher’s least significant difference for comparisons among groups. Values of *p* < 0.05 were considered statistically significant.

## Results

### Generation of iPSCs

To generate iPSCs, dermal fibroblasts of a healthy donor were nucleofected with episomal vectors expressed reprogramming factors. Colonies similar to those of human pluripotent cells formed beginning from day 15 after nucleofection. These colonies were cultivated to generate stable cell lines. Seventeen cell lines were derived, and four of these cell lines (m34Sk3, m34Sk4, m34Sk10, and m34Sk13) were selected for further cultivation and characterization.

The cell lines obtained expressed a number of pluripotency markers, such as alkaline phosphatase (Fig. 1A), transcription factors (OCT4 and NANOG), and surface antigens (SSEA-4, TRA-1-60, and TRA-1-81) (Fig. 1B). The cell lines underwent spontaneous differentiation through embryoid body formation The differentiated derivatives obtained during spontaneous differentiation expressed markers of three germ layers: ectoderm (beta-III-tubulin), endoderm (cytokeratin 18), and mesoderm (CD90, smooth muscle α-actin) (Fig. 1C). Expression of pluripotency markers and capacity to be differentiated into three germ layer derivatives confirmed the pluripotent state of the cell lines obtained.

**Fig. 1.**
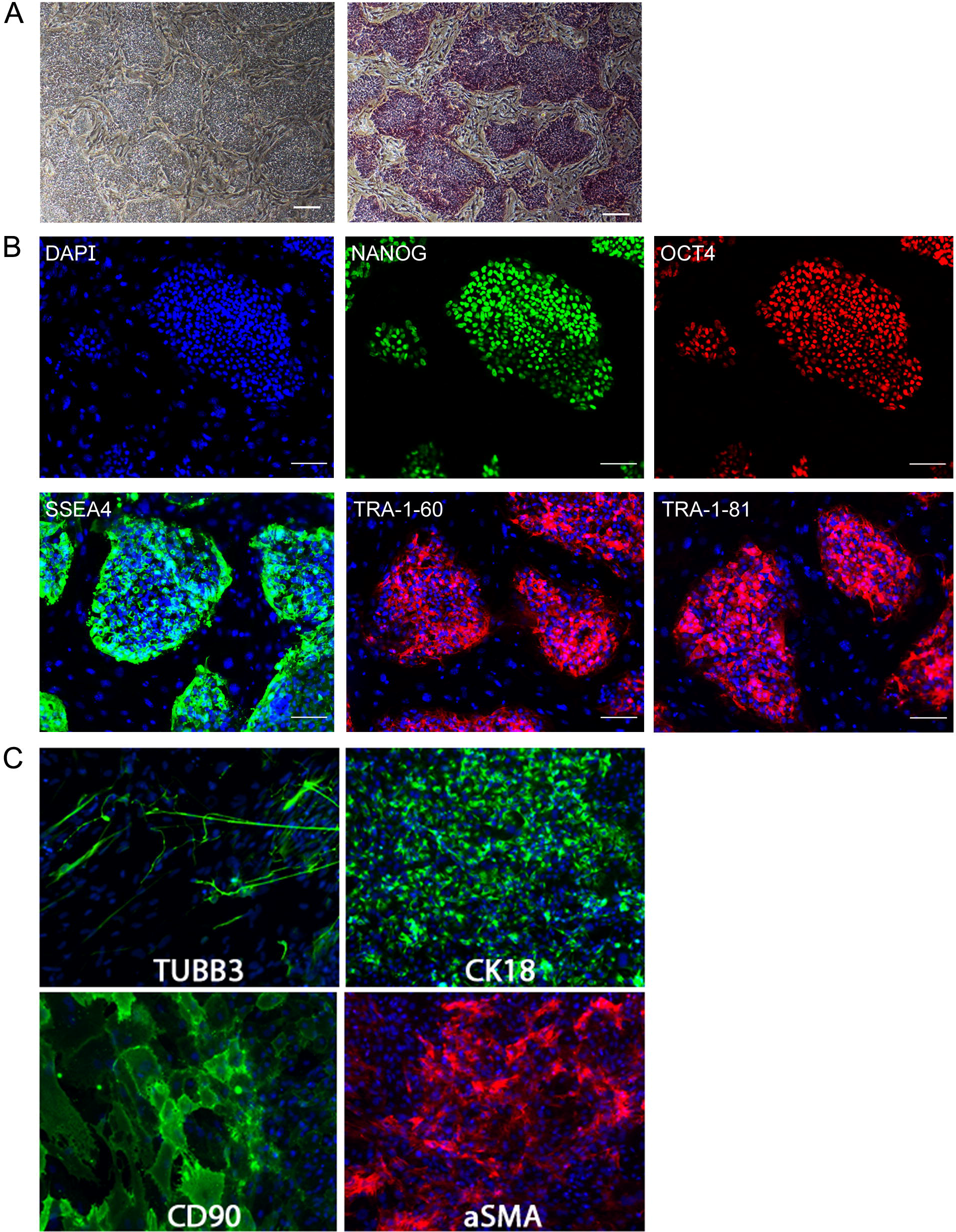
Characterization of iPSCs obtained from a healthy donor. A) Morphology of the cell lines generated by reprogramming of fibroblasts to a pluripotent state (on the left) and the expression of alkaline phosphatase in the cell lines (on the right). Scale bar: 200 μM. B) The expression of NANOG and OCT4 transcription factors and SSEA-4, TRA-1-60, and TRA-1-81 surface antigens. The nuclei were stained with DAPI. Scale bar: 100 μM C) Spontaneous differentiation pattern. The expression of three germ layer markers in differentiated derivatives: beta-III-tubulin (ectoderm), cytokeratin 18 (endoderm), smooth muscle α actin, and CD90 (mesoderm). The nuclei were stained with DAPI Scale bar: 100 μm. C) Spontaneous differentiation pattern. The expression of three germ layer markers in differentiated derivatives: beta-III-tubulin (ectoderm), cytokeratin 18 (endoderm), smooth muscle α-actin, and CD90 (mesoderm). The nuclei were stained with DAPI

### Directed differentiation of iPSCs into cardiomyocytes

The method based on WNT/β catenin activation by inhibition of the GSK3 protein kinase with CHIR99021, followed by WNT repression (with IWP2) [17] was applied to induce iPSC differentiation into cardiomyocytes. One of the iPSC lines, m34Sk3, was differentiated into cardiomyocytes, with the first spontaneous contractions observed on day 9. The analysis of single differentiated cells revealed the expression of sarcomeric proteins (cardiac troponin T, ventricular form of β-myosin regulatory light chain, and β yosin heavy chain) and cardiomyocyte transcription factor NKX2.5 in a proportion of the cells (Fig. 2).

**Fig. 2.**
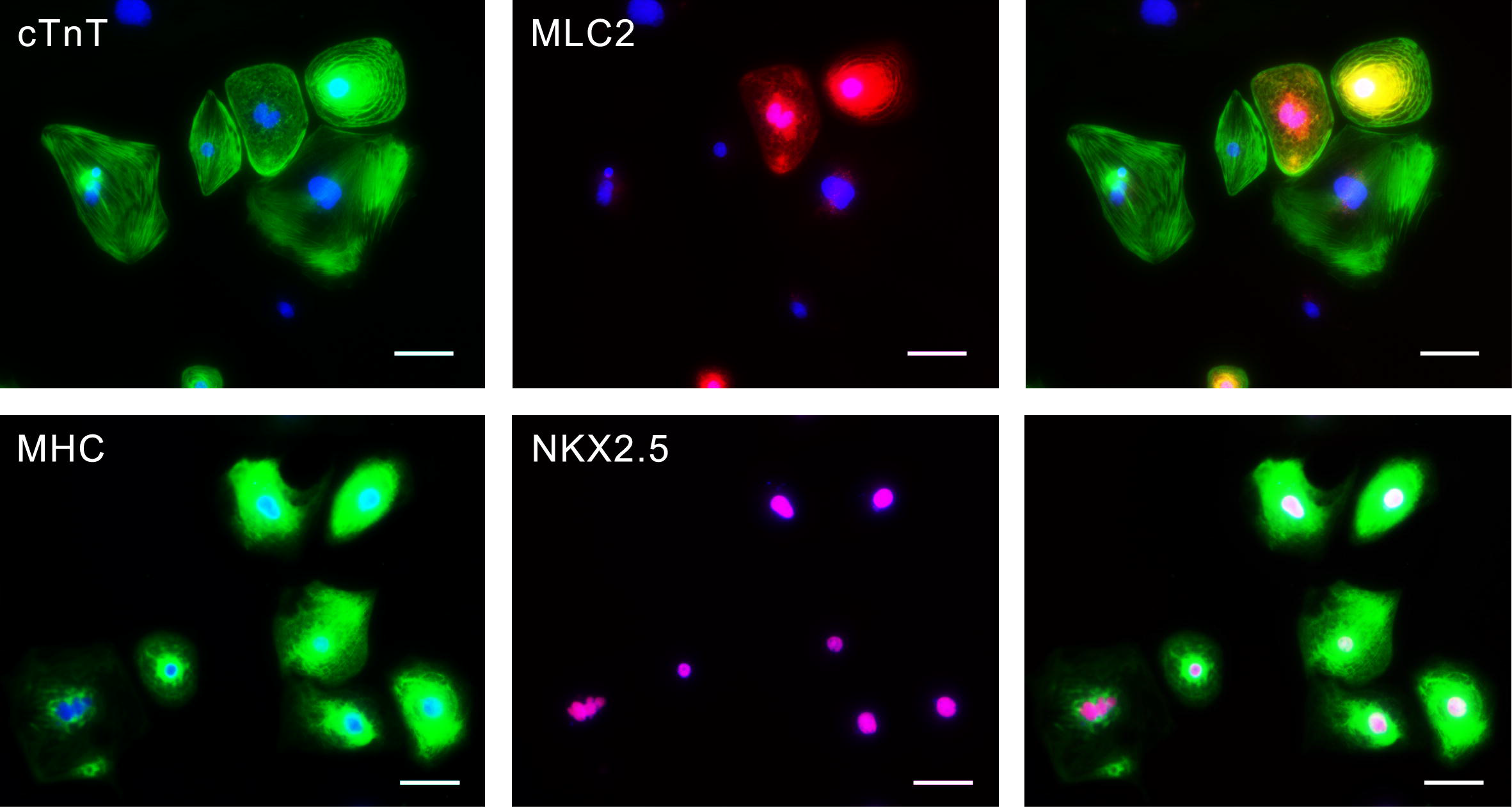
Directed differentiation of iPSCs into cardiomyocytes. The expression of sarcomeric proteins — cardiac troponin T (cTnT), β yosin regulatory light chain (MLC2), β yosin heavy chain (MHC), and cardiomyocyte transcription factor NKX2.5 in differentiated cells of the m34Sk3 line. The nuclei were stained with DAPI. Scale bar: 50 μm

### Optical mapping of human iPSC-derived cardiomyocytes

In the first set of experiments, monolayers of human iPSC-derived cardiomyocytes were used to evaluate excitation wave propagation under erythromycin concentrations of 15, 30, and 45 μM. The results revealed that the CV was changed insignificantly (the maximal decrease 12±9% under 30 μM) under these concentrations (Fig. 3A) and that the maximum capture rate decreased up to 28±12% (Fig. 3B, *p* < 0.05). The latter effect was irreversible after washout of erythromycin.

**Fig. 3.**
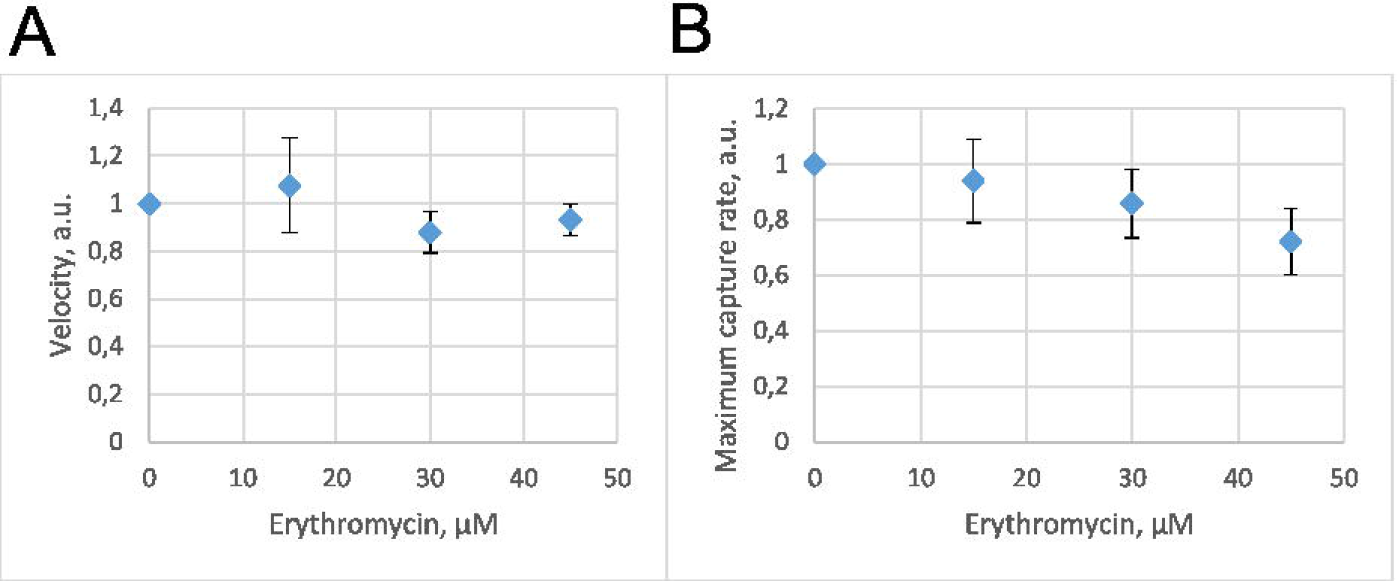
The effect of erythromycin on the conduction velocity (CV) and maximum capture rate (MCR) in the human iPSC-derived cardiomyocyte monolayer. The CV (A) and MCR (B) at erythromycin concentrations of 15, 30, and 45 μM (*p* < 0.05). The CV and MCR were measured 5–15 min after the addition of erythromycin solution. The frequency was increased from 1 Hz to 3.33 Hz in increments of ≤ 0.5 Hz

Second, we evaluated the formation of re-entry on a standard linear obstacle situated perpendicular to the direction of excitation wave propagation. The frequency of external impulses was increased from 1 Hz to 3.33 Hz in increments of ≤ 0.5 Hz. Figure 4 shows the activation maps of the human iPSC-derived cardiomyocyte monolayer with the obstacle at a stimulation frequency of 2.5 Hz in the control (4A) and when exposed to an erythromycin concentration of 15 μM (4B).

**Fig. 4.**
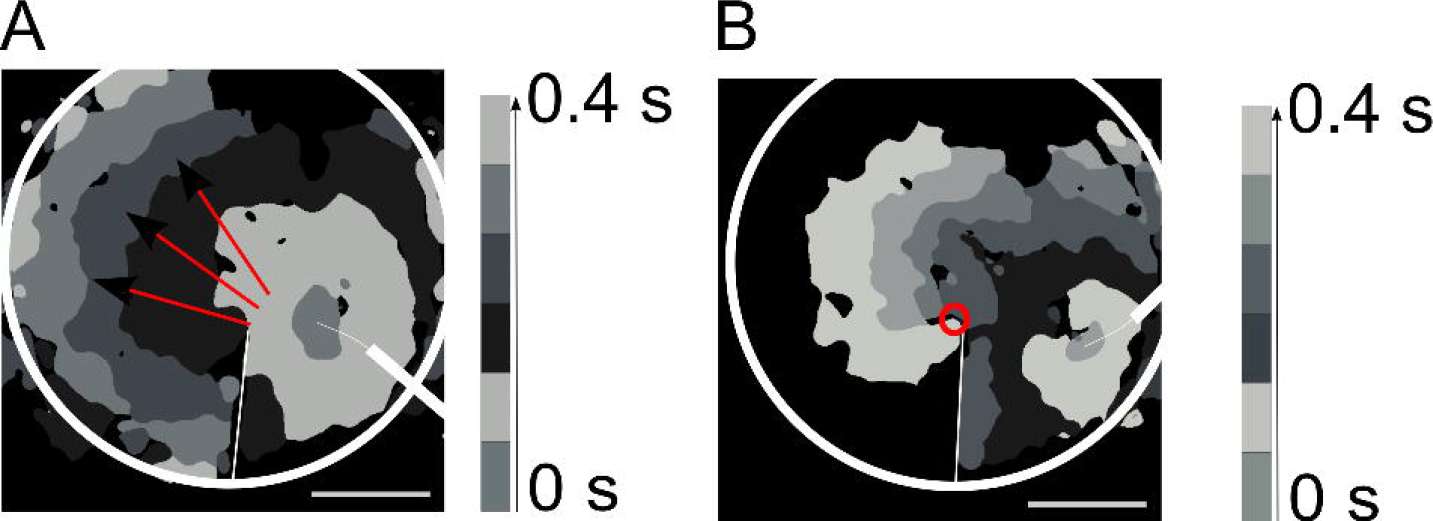
Activation maps of the human iPSC-derived cardiomyocyte monolayer with the standard linear obstacle located perpendicular to the excitation wave propagation front. The stimulation frequency was 2.5 Hz. The white circle corresponds to the borders of the sample, and the white line corresponds to the location of the obstacle. Scale bar: 3.48 mm. A) Control, normal propagation on the obstacle. The red arrows indicate the direction of excitation wave propagation. B) Erythromycin (15 μM showing re-entry formation on the sharp end of the obstacle. The red circle indicates the core of re-entry

Figure 5 shows diagrams of the probability of re-entry occurrence under erythromycin concentrations of 15, 30, and 45 μM as a function of the stimulation frequency (blue bars). With an increase in the concentration of erythromycin, re-entry occurred at lower stimulation frequencies. The orange bars indicate cases where the monolayer did not capture the external impulses.

**Fig. 5.**
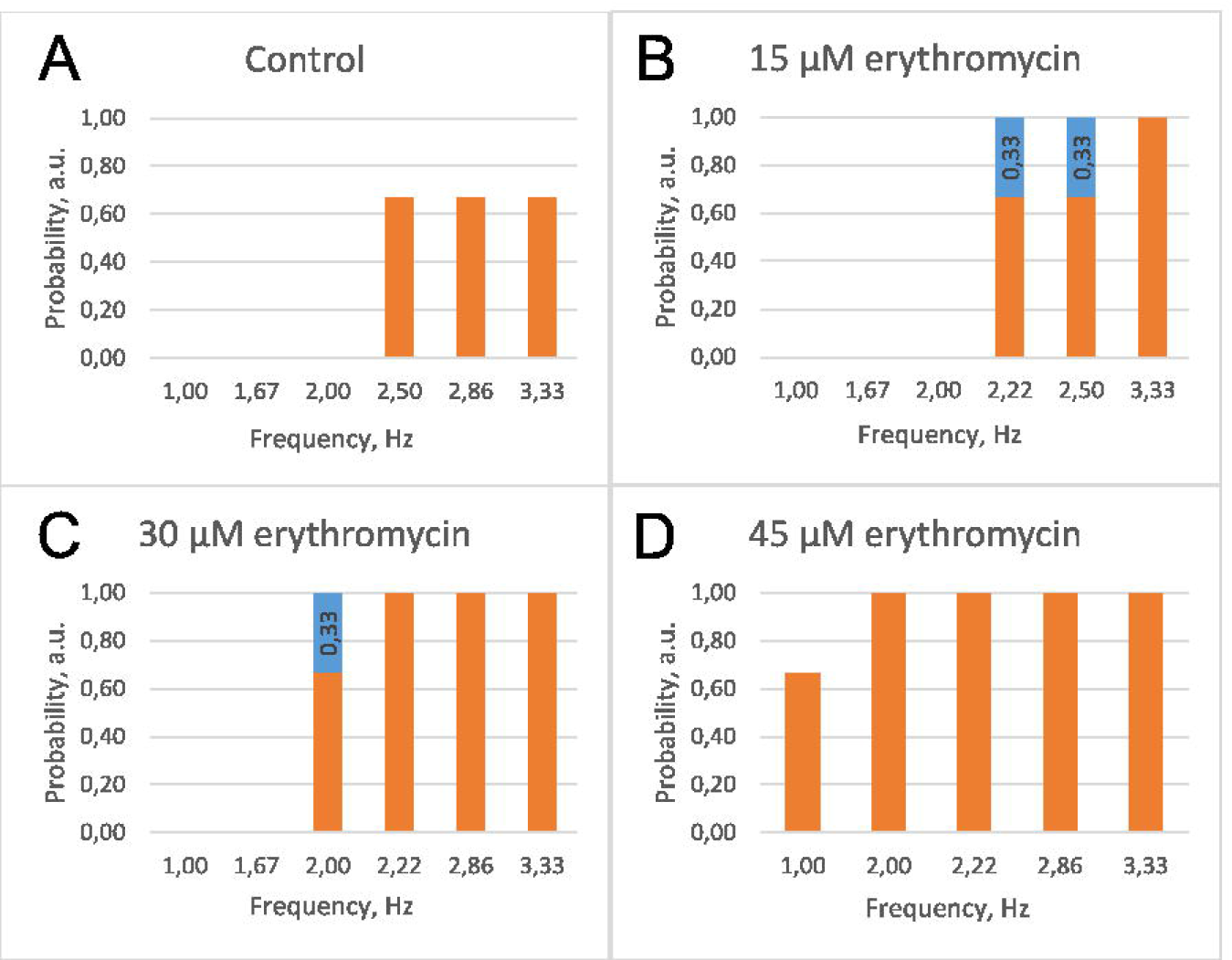
The effect of erythromycin on re-entry formation in the human iPSC-derived cardiomyocyte monolayer. The orange bars indicate cases where the monolayer did not capture the external impulses. The blue bars indicate re-entry occurrence. Re-entry occurred at erythromycin concentrations of 15 and 30 μM

### Optical mapping of NRVMs

In the second set of experiments, excitation wave propagation in NRVM monolayers under erythromycin concentrations of 15, 30, and 45 μM was evaluated. The MCR was stable (a decrease of less than 9±8%) under these concentrations (Fig. 6A). Re-entry did not occur at all the concentrations of erythromycin (15, 30, and 45 μM). Figure 6C demonstrates the successful excitation wave propagation in NRVM monolayer with the standard linear obstacle under 15 μM erythromycin at stimulation frequency of 3.33 Hz.

**Fig. 6.**
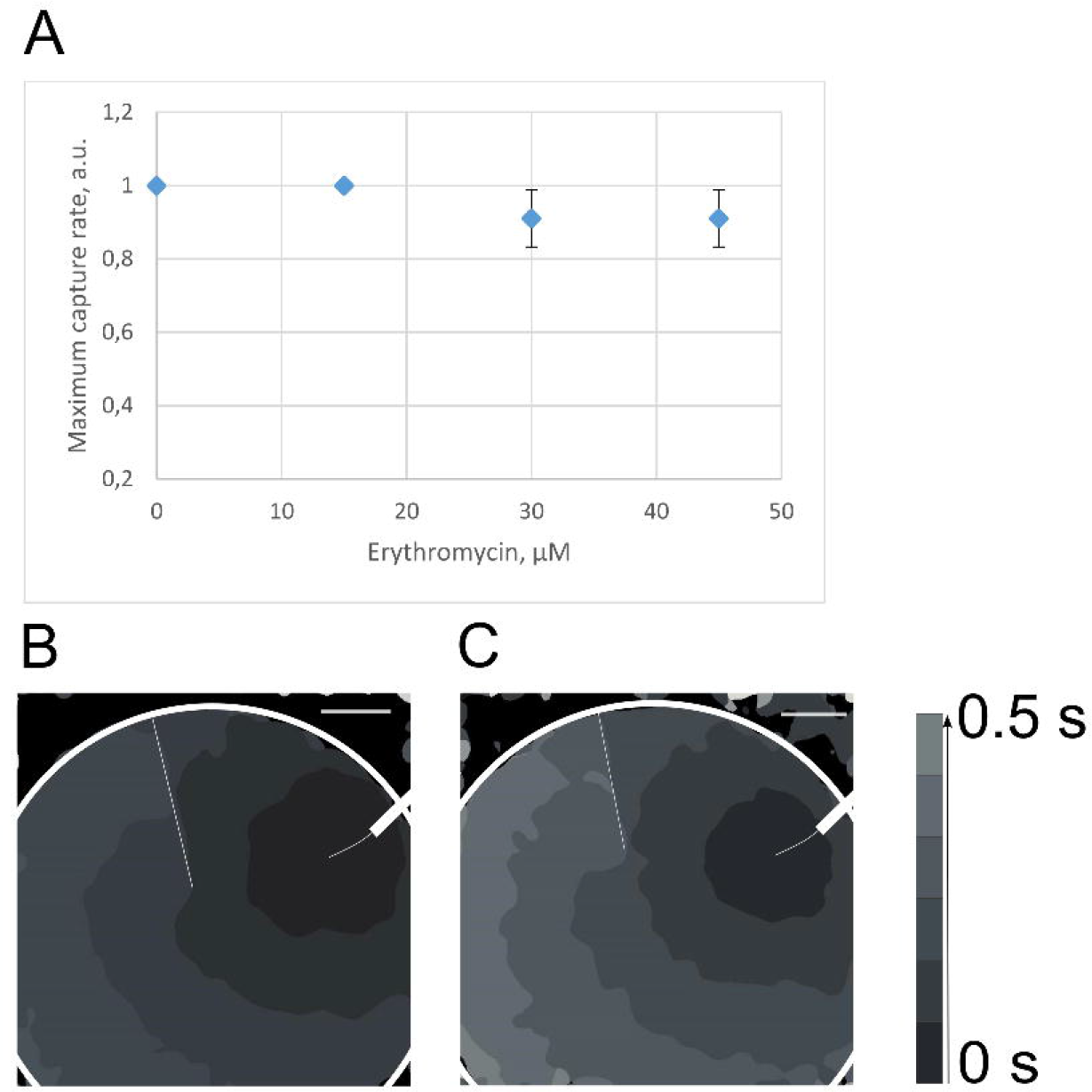
The effect of erythromycin on excitation wave propagation in the NRVM monolayer. A) The maximum capture rate (MCR) at erythromycin concentrations of 15, 30, and 45 μM. The MCR was measured 5–15 min after the addition of erythromycin solution. The MCR was increased from 1 Hz to 3.33 Hz in increments of ≤ 0.5 Hz. B,C) Activation maps of the NRVM monolayer with the standard linear obstacle located perpendicular to the excitation wave propagation front. The white circle corresponds to the borders of the sample, and the white line corresponds to the location of the obstacle. Scale bar: 1 mm. B) Control, with a stimulation frequency of 1 Hz. C) Erythromycin at a concentration of 15 μM and stimulation frequency of 3.33 Hz. Re-entry did not occur at a concentration of 15 μM or at higher concentrations (30 and 45 μM) of erythromycin

## Discussion

In the last decade [16,17] in pharmacology, cell assays have been widely used for drug testing. Today, cellular assays can exploit cell reprogramming [10, 20]. Using this technology, adult cells of a living organism can be reprogrammed into a pluripotent state. Following further differentiation of these cells, human cells with specific properties can be obtained *in vitro* [19–21].

In standard cardiotoxicity assays, a chemical substance is used to inhibit hERG channels to identify possible side effects namely the occurrence of life-threatening arrhythmias, e.g. TdP, associated with the substance [24, 25]. Few studies show the effect of arrhythmogenic drugs on arrhythmic beating, changes in beat rate [2] and FPD [14] using monolayers of human iPSC-derived cardiomyocytes (iCell®). Shaheen et al. generated human iPSC-derived cardiac cell sheets, expressing the genetically encoded voltage indicator ArcLight for evaluation of conduction slowing and APD prolongation under chemical compounds [15]. In addition, the developing systems [26] of patient-specific human iPSC-derived cardiomyocytes consider a single cell level. A shortcoming of existing methods of drug testing for cardiotoxicity based on iPSCs is a small consideration of the drug effect on intercellular contacts and conduction of excitation in multicellular, syncytium-forming systems.

To enhance the reliability of cardiotoxicity testing systems, we propose a method for determining the arrhythmogenicity of the substance based on evaluating re-entry formation on a standard linear obstacle in cardiomyocyte monolayer by optical mapping. We validated the proposed method in two model systems: a monolayer of human iPSC-derived cardiomyocytes from healthy donor and a monolayer of NRVM. As a test substance, erythromycin, a representative macrolide antibiotic that prolongs the QT interval and potentially induces TdP, was used.

In the experiments on the dose dependence of the conduction velocity (CV) and maximum capture rate (MCR) of the excitation wave in the monolayers of human iPSC-derived cardiomyocytes the CV varied insignificantly (up to 12 ± 9%) at erythromycin concentrations of 15–45 μM as compared with that of the control, whereas the MCR fell substantially (to 28 ± 12%) at these concentrations. As the MCR of stimulation corresponds to the total duration of the action potential and the period of refractoriness, the decrease in this parameter points to prolongation of the sum of the above characteristics and indirectly confirms the action of erythromycin on the I_Kr_ current. This potassium current is present in human cells but absent in rat cells. In the experiments on MCR in the monolayer of rat cardiomyocytes, no similar effects were obtained, and the MCR changed insignificantly when exposed to erythromycin at concentrations of 15–45 μM (Fig. 6A). These findings confirm the results of a where re-entry did not occur at a coprevious study on the effect of erythromycin on the I_Kr_ current [5-8] and the lack of any effect on the I_Ks_ current [6].

Although the active concentration of erythromycin is 30 μM [2], erythromycin-induced effects have been observed in the range of 1.5 μM up to 40 μM and even 100μM [12]. In the present study a noticeable cardiotoxic effect (MCR decrease) was observed at concentrations of 15–45 μM.

In the experiments on human iPSC-derived cardiomyocyte monolayers with a standard linear obstacle, re-entry formation was observed on a sharp end of the obstacle at concentrations of 15 and 30 μM where as re-entry did not occur at a concentration of 45 μM. These findings provide evidence for a “window” at which concentrations of the drug lead to re-entry formation and in which erythromycin should be administered with extreme caution. For example, the clinically effective concentration of erythromycin (30 μM) is clearly dangerous. Thus, small doses of the drug, with a final concentration not exceeding 15 μM are likely advisable in patients with an increased heart rate (i.e., higher than 100 beats/min) to avoid side effects manifesting at erythromycin blood concentrations between 15 μM and 30 μM.

## Conclusions

The proposed test method, which is based on a comparison of MCR in human iPSC-derived cardiomyocyte and NRVM monolayers by optical mapping, provides indirect confirmation of the effect of erythromycin on the delayed rectifier current (I_Kr_ and I_Ks_). The tissue model of the monolayer of iPSC-derived cardiomyocytes pointed to the presence of a concentrations window (15–30 μM) at which the probability of the occurrence of re-entry increased. Using a standard linear obstacle, this study demonstrates for the first time the mechanism underlying the occurrence of re-entry using a simplified model of a cardiomyocytes monolayer after the application of 15, 30, 45 μM erythromycin.

## Ethical approval

All procedures performed in studies involving human participants were in accordance with the ethical standards of the institutional and/or national research committee and with the 1964 Helsinki declaration and its later amendments or comparable ethical standards.

All applicable international, national, and/or institutional guidelines for the care and use of animals were followed.

## Acknowledgments

This study was funded by the Russian Ministry of Education and Science of the Russian Federation grant (state task) 6.9906.2017/BCh.

## References

1. Gibson, J. K., Yue, Y., Bronson, J., Palmer, C., & Numann, R. (2014). Human stem cell-derived cardiomyocytes detect drug-mediated changes in action potentials and ion currents. Journal of Pharmacological and Toxicological Methods, 70(3), 255–267. doi:10.1016/J.VASCN.2014.09.005

2. Guo, L., Coyle, L., Abrams, R. M. C., Kemper, R., Chiao, E. T., & Kolaja, K. L. (2013). Refining the Human iPSC-Cardiomyocyte Arrhythmic Risk Assessment Model. Toxicological Sciences, 136(2), 581–594. doi:10.1093/toxsci/kft205

3. Ray, W. A., Murray, K. T., Meredith, S., Narasimhulu, S. S., Hall, K., & Stein, C. M. (2004). Oral Erythromycin and the Risk of Sudden Death from Cardiac Causes. New England Journal of Medicine, 351(11), 1089–1096. doi:10.1056/NEJMoa040582

4. Itoh, H., Sakaguchi, T., Ding, W.-G., Watanabe, E., Watanabe, I., Nishio, Y., … Horie, M. (2009). Latent Genetic Backgrounds and Molecular Pathogenesis in Drug-Induced Long-QT Syndrome. Circulation: Arrhythmia and Electrophysiology, 2(5), 511–523. doi:10.1161/CIRCEP.109.862649

5. Duncan, R. S., Ridley, J. M., Dempsey, C. E., Leishman, D. J., Leaney, J. L., Hancox, J. C., & Witchel, H. J. (2006). Erythromycin block of the HERG K+ channel: Accessibility to F656 and Y652. Biochemical and Biophysical Research Communications, 341(2), 500–506. doi:10.1016/J.BBRC.2006.01.008

6. Antzelevitch, C., Sun, Z.-Q., Zhang, Z.-Q., & Yan, G.-X. (1996). Cellular and Ionic Mechanisms Underlying Erythromycin-Induced Long QT Intervals and Torsade de Pointes. Journal of the American College of Cardiology, 28(7), 1836–1848. doi:10.1016/S0735-1097(96)00377-4

7. Smits, J. P., Blom, M. T., Wilde, A. A., & Tan, H. L. (2008). Cardiac sodium channels and inherited electrophysiologic disorders: a pharmacogenetic overview. Expert Opinion on Pharmacotherapy, 9(4), 537–549. doi:10.1517/14656566.9.4.537

8. Lazzerini, P. E., Yue, Y., Srivastava, U., Fabris, F., Capecchi, P. L., Bertolozzi, I., … Boutjdir, M. (2016). Arrhythmogenicity of Anti-Ro/SSA Antibodies in Patients With Torsades de Pointes. Circulation: Arrhythmia and Electrophysiology, 9(4), e003419. doi:10.1161/CIRCEP.115.003419

9. Schulz, M., & Schmoldt, A. (2003). Therapeutic and toxic blood concentrations of more than 800 drugs 8 and other xenobiotics. Die Pharmazie, 58(7), 447–74. Retrieved from http://www.ncbi.nlm.nih.gov/pubmed/12889529

10. Kuusela, J., Kujala, V. J., Kiviaho, A., Ojala, M., Swan, H., Kontula, K., & Aalto-Setälä, K. (2016). Effects of cardioactive drugs on human induced pluripotent stem cell derived long QT syndrome cardiomyocytes. SpringerPlus, 5(1), 234. doi:10.1186/s40064-016-1889-y

11. Stanat, S. J. C., Carlton, C. G., Crumb Jr, W. J., Agrawal, K. C., & Clarkson, C. W. (2003). Characterization of the inhibitory effects of erythromycin and clarithromycin on the HERG potassium channel. Molecular and Cellular Biochemistry, 254(1/2), 1–7. doi:10.1023/A:1027309703313

12. Guo, J., Zhan, S., Lees-Miller, J. P., Teng, G., & Duff, H. J. (2005). Exaggerated block of hERG (KCNH2) and prolongation of action potential duration by erythromycin at temperatures between 37°C and 42°C. Heart Rhythm, 2(8), 860–866. doi:10.1016/J.HRTHM.2005.04.029

13. Liu, Y., Xia, T., Wei, J., Liu, Q., & Li, X. (2017). Micropatterned co-culture of cardiac myocytes on fibrous scaffolds for predictive screening of drug cardiotoxicities. Nanoscale, 9(15), 4950–4962. doi:10.1039/C7NR00001D

14. Ando, H., Yoshinaga, T., Yamamoto, W., Asakura, K., Uda, T., Taniguchi, T., … Sekino, Y. (2017). A new paradigm for drug-induced torsadogenic risk assessment using human iPS cell-derived cardiomyocytes. Journal of Pharmacological and Toxicological Methods, 84, 111–127. doi:10.1016/j.vascn.2016.12.003

15. Shaheen, N., Shiti, A., Huber, I., Shinnawi, R., Arbel, G., Gepstein, A., … Gepstein, L. (2018). Human Induced Pluripotent Stem Cell-Derived Cardiac Cell Sheets Expressing Genetically Encoded Voltage Indicator for Pharmacological and Arrhythmia Studies. Stem Cell Reports, 10(6), 1879–1894. doi:10.1016/j.stemcr.2018.04.006

16. Medvedev, S. P., Grigor’eva, E. V., Shevchenko, A. I., Malakhova, A. A., Dementyeva, E. V., Shilov, A. A., … Zakian, S. M. (2011). Human Induced Pluripotent Stem Cells Derived from Fetal Neural Stem Cells Successfully Undergo Directed Differentiation into Cartilage. Stem Cells and Development, 20(6), 1099–1112. doi:10.1089/scd.2010.0249

17. Burridge, P. W., Matsa, E., Shukla, P., Lin, Z. C., Churko, J. M., Ebert, A. D., … Wu, J. C. (2014). Chemically defined generation of human cardiomyocytes. Nature Methods, 11(8), 855–860. doi:10.1038/nmeth.2999

18. Astashkina, A., Mann, B., & Grainger, D. W. (2012). A critical evaluation of in vitro cell culture models for high-throughput drug screening and toxicity. Pharmacology & Therapeutics, 134(1), 82–106. doi:10.1016/J.PHARMTHERA.2012.01.001

19. Kepp, O., Galluzzi, L., Lipinski, M., Yuan, J., & Kroemer, G. (2011). Cell death assays for drug discovery. Nature Reviews Drug Discovery, 10(3), 221–237. doi:10.1038/nrd3373

20. Scott, C. W., Peters, M. F., & Dragan, Y. P. (2013). Human induced pluripotent stem cells and their use in drug discovery for toxicity testing. Toxicology Letters, 219(1), 49–58. doi:10.1016/J.TOXLET.2013.02.020

21. Grskovic, M., Javaherian, A., Strulovici, B., & Daley, G. Q. (2011). Induced pluripotent stem cells — opportunities for disease modelling and drug discovery. Nature Reviews Drug Discovery, 10(12), 915. doi:10.1038/nrd35779

22. Avior, Y., Sagi, I., & Benvenisty, N. (2016). Pluripotent stem cells in disease modelling and drug discovery. Nature Reviews Molecular Cell Biology, 17(3), 170–182. doi:10.1038/nrm.2015.27

23. Itzhaki, I., Maizels, L., Huber, I., Zwi-Dantsis, L., Caspi, O., Winterstern, A., … Gepstein, L. (2011). Modelling the long QT syndrome with induced pluripotent stem cells. Nature, 471(7337), 225–229. doi:10.1038/nature09747

24. Braam, S. R., Tertoolen, L., van de Stolpe, A., Meyer, T., Passier, R., & Mummery, C. L. (2010). Prediction of drug-induced cardiotoxicity using human embryonic stem cell-derived cardiomyocytes. Stem Cell Research, 4(2), 107–116. doi:10.1016/J.SCR.2009.11.004

25. Tanaka, T., Tohyama, S., Murata, M., Nomura, F., Kaneko, T., Chen, H., … Fukuda, K. (2009). In vitro pharmacologic testing using human induced pluripotent stem cell-derived cardiomyocytes. Biochemical and Biophysical Research Communications, 385(4), 497–502. doi:10.1016/J.BBRC.2009.05.073

26. Liang, P., Lan, F., Lee, A. S., Gong, T., Sanchez-Freire, V., Wang, Y., … Wu, J. C. (2013). Drug Screening Using a Library of Human Induced Pluripotent Stem Cell–Derived Cardiomyocytes Reveals Disease-Specific Patterns of Cardiotoxicity. Circulation, 127(16), 1677–1691. doi:10.1161/CIRCULATIONAHA.113.001883

